# The Impact of Multiple Sequence Alignment Error on Phylogenetic Estimation under Variable-Across-Phylogeny Substitution Models

**DOI:** 10.1101/2023.01.24.525451

**Authors:** Rei Doko, Kevin J. Liu

## Abstract

In many different species, it has been observed that nucleotide compositions are not identical on the genic and even genomic scale. This observation contradicts a commonly held assumption in most maximum likelihood based phylogenetic estimation methods - that the process governing DNA evolution is identical across lineages. We show that when DNA evolution is nonhomogeneous, topological estimation and continuous parameter estimation are impacted both by alignment quality and model misspecification due to the homogeneity-across-lineages assumption.

## 1 Introduction

Nucleotide composition biases can be found in the genomes of a variety of organisms, such as grasses [**?**], insects [**?**], and birds [**?**]. Knowing when and where these compositional biases arise in the evolutionary history of these organisms is of interest since G+C bias is hypothesized to have significance in biological processes. Computational methods can be applied to more widely available genomic data to provide a better idea of this history. Molecular phylogenetics is used to reconstruct evolutionary relationships between organisms using biomolecular sequence data such as DNA.

Maximum likelihood based phylogenetic estimation uses a stochastic model of sequence evolution to evaluate the probability of a tree topology given the observed sequence data. A common simplifying assumption is that the base composition and relative substitution rates are identical throughout the tree. However, the observation that nucleotide composition can vary across lineages demonstrates that these assumptions do not hold in biological data. Nonhomogeneous substitution models relax this assumption and allow for base composition and relative substitution rates to vary across the phylogeny, and have been implemented before in nhPhyML [**?**] and PAML [**?**].

Alignment error has been shown to impact downstream phylogenetic inference and estimation [**?**]. However, how alignment quality impacts estimation when sequence evolution is nonhomogeneous is not well studied. While it is likely that alignment quality will have an impact on estimation in this more complicated model, the question of what does this mean for empirical data remains. For example, how are estimates of base composition and substitution rates when using nonhomogeneous substitution models for maximum likelihood estimation? Furthermore, do more sophisticated models of DNA evolution accounting for nonhomogeneity improve estimates?

## 2 Materials and Methods

The objective of this study is to characterize the effect of alignment quality and model misspecification in the problem of phylogenetic estimation when evolution is nonhomogeneous and nonstationary.

### Data Availability Statement

Data and scripts used are available at https://gitlab.msu.edu/liulab/nonhomogeneous-substitution-model-study-data-scripts.

### 2.1 Methods for MSA and phylogenetic estimation

#### Preliminaries

Let *T* = (*V, E*) be a rooted tree with labeled leaves *X* ⊂ *V* and root *ρ* ∈ *V*. Each edge *e* = (*u, v*) ∈ *E* where *u,v* ∈ V has a length d(e). An edge (*u,v*) is a leaf edge if either *u* or *v* is a leaf, otherwise it is an internal edge. Deleting an edge *e* from a tree *T* gives two subtrees *T*_1_ = (*V*_1_, *E*_1_) and *T*_2_ = (*V*_2_, *E*_2_). The vertex sets *V*_1_ and *V*_2_ are disjoint, and *V*_1_ ∪ *V*_2_ = *V*. The same can be said for their respective leaf sets, so {*X*_1_, *X*_2_} is a bipartition of *X*. Let this be denoted as *b*(*e*) = {*X*_1_, *X*_2_}.

#### Multiple sequence alignment

There are a variety of multiple sequence alignment methods available. For this study, we selected a range of commonly used methods. We aligned simulated and empirical datasets using MAFFT [**?**] version 7.475, MUSCLE [**?**] version 5.0.1428, Clustal Omega [**?**] version 1.2.4, Clustal W [**?**] version 2.1, and FSA version [**?**] 1.15.9. Each method was run using their respective default settings.

#### Phylogenetic estimation

We use the general time reversible (GTR) model for phylogenetic estimation. The GTR model specifies that there are separate base frequencies *π_T_, π_C_, π_A_, π_G_* which sum to 1, as well as rate parameters *a, b, c, d, e, f*. We use the same conventions as used by **?**. That is to say, *a* corresponds to *T* ↔ *C, b* corresponds to *T* ↔ *A, c* to *T* ↔ *G, d* to *C* ↔ *A, e* to *C* ↔ *G*, and *f* to *G* ↔ *A*. f is fixed to 1 and the remaining rate parameters are relative to *f*. The rate matrix *Q* is as follows:

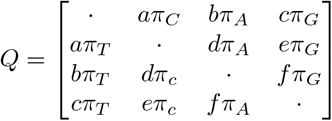

With the diagonals set to *Q_ii_* = ∑_*i*≠*j*_ *Q_ij_*. The transition probability matrix is given by *P*(*t*) = exp(−*Qt*) and is used to calculate likelihoods for a phylogenetic tree. Typically in phylogenetic estimation using Markov models of substitution, the rate matrix is assumed to be constant over the whole tree. We refer to models under this assumption as homogeneous, or having no shifts.

However, nucleotide composition biases have been observed in biological data. To account for rate and composition differences across lineages, each edge e has an associated set of parameters *θ*(*e*) that define the rate matrix for that edge. We use a GTR model for the branch models. The traditional homogeneous model is the case where *θ*(*e*) is fixed, i.e. *θ*(*e_i_*) = *θ*(*e_j_*) for all *e_i_, e_j_* ∈ *E*. For heterogeneous models, we considered two different classes: which we refer to as single-shift and all-shift. For all-shift, *θ*(*e*) is independent for each edge. For single-shift, there are exactly two sets of parameters, *θ*_shift_ and *θ*_background_ and some restrictions on which edges they apply to. There is a shift edge, *e*_shift_ ∈ *E*, and all edges descending from it all have *θ*(*e*) = *θ*_shift_. Any remaining edges are *θ*_background_. We will also refer to homogeneous models as no-shift interchangeably. In nonhomogeneous models, rooting can impact likelihood values since these models are not time-reversible, so rooted trees are used.

RAxML [**?**] version 8.2.12 was used to perform maximum likelihood estimation under a homogeneous GTR model. PAML [**?**] version 4.9j was used to perform maximum likelihood estimation under fixed tree topologies using a branch model. Since PAML does not support tree search under nonhomogeneous models, we wrote a wrapper script to perform local tree topology search, using PAML to evaluate the log likelihood of a topology.

#### Single-shift search

While a fully nonhomogeneous branch model can account for very general nucleotide substitution processes, it is highly parameterized. Also, it approaches the no common mechanism model, which is known to be statistically inconsistent [**?**]. In single-shift, there are exactly two separate sets of substitution model parameters estimated, and instead is search for a placement of the different rate matrix on the phylogeny. We use a bruteforce approach of local search to determine which assignment of the shift model maximizes likelihood.

#### Performance assessments

We use Robinson-Foulds distance [**?**] to assess topological difference between trees. Let 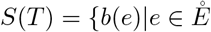, *e* is an internal edge}. The Robinson-Foulds distance between two unrooted trees *T* and *T*′ is the symmetric difference of *S*(*T*) and *S*(*T*′). A way to extend RF distance to rooted trees is to consider the bipartition representation for labeled nodes (i.e. *X*∪{*ρ*}). So for the two subtrees *T*_1_, *T*_2_ induced by deleting an edge *e* ∈ *E*, if *ρ* ∈ *V*_1_ then the edge representation becomes *b*′(*e*) = {*X*_1_ ∪ {*ρ*}, *X*_2_} and vice versa if *ρ* ∈ *V*_2_. For identifying root placement, we say two trees *T* and *T*′ have identical roots, *ρ* and *ρ*′ respectively, if the leaf sets of the subtrees induced by deleting the respective root nodes are identical.

We take the L1 norm of the relative errors for substitution model parameters to assess model parameter estimation performance in the simulation study. For the base frequencies, this would be 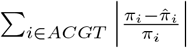.

To assess how well the shift subtree is being predicted in the single-shift model, we use the size of the maximum agreement subtree (MAST) between the true and estimated shift subtrees. The MAST problem is to find a subtree given a set of trees 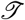 with the largest subset of leaves that also agrees with all the trees in 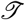.

For evaluating alignment quality, we use sum-of-pairs false positive and false negative rates, denoted SP-FP and SP-FN respectively. SP-FP is calculated as the proportion of homologies in the estimated alignment and not in the true alignment. SP-FN is the same, but the other way around.

### 2.2 Simulation study

#### Model tree generation

Model trees were sampled using INDELible [**?**] under a birth-death process. Non-ultrametricity was introduced using a procedure described in **?** with deviation factor *c* = 2.

1. Generate a rooted model tree using INDELible with the default settings
2. For every branch:
  2.1 Choose *x* ~ *U*(– ln(2),ln(2))
  2.2 Scale the branch length by exp(*x*)
3. Let *L* be the maximum root-to-tip distance for the tree and *H* be the desired height.
4. Scale each branch length by *H/L*
5. Select a subtree to evolve under the shifted substitution model containing as close to half of the leaves.

**Table 1:** GTR model parameters used for evolving sequences in the simulation study.

| Parameter | T | C | A | G | C↔T | A↔T | G↔T | A↔C | C↔G | A↔G |
| --- | --- | --- | --- | --- | --- | --- | --- | --- | --- | --- |
| Shift | 0.216 | 0.237 | 0.317 | 0.230 | 5.847 | 3.186 | 1.214 | 3.437 | 1.307 | 1.0 |
| Background | 0.183 | 0.226 | 0.058 | 0.534 | 1.505 | 0.367 | 0.141 | 0.412 | 0.094 | 1.0 |

**Figure 1:**
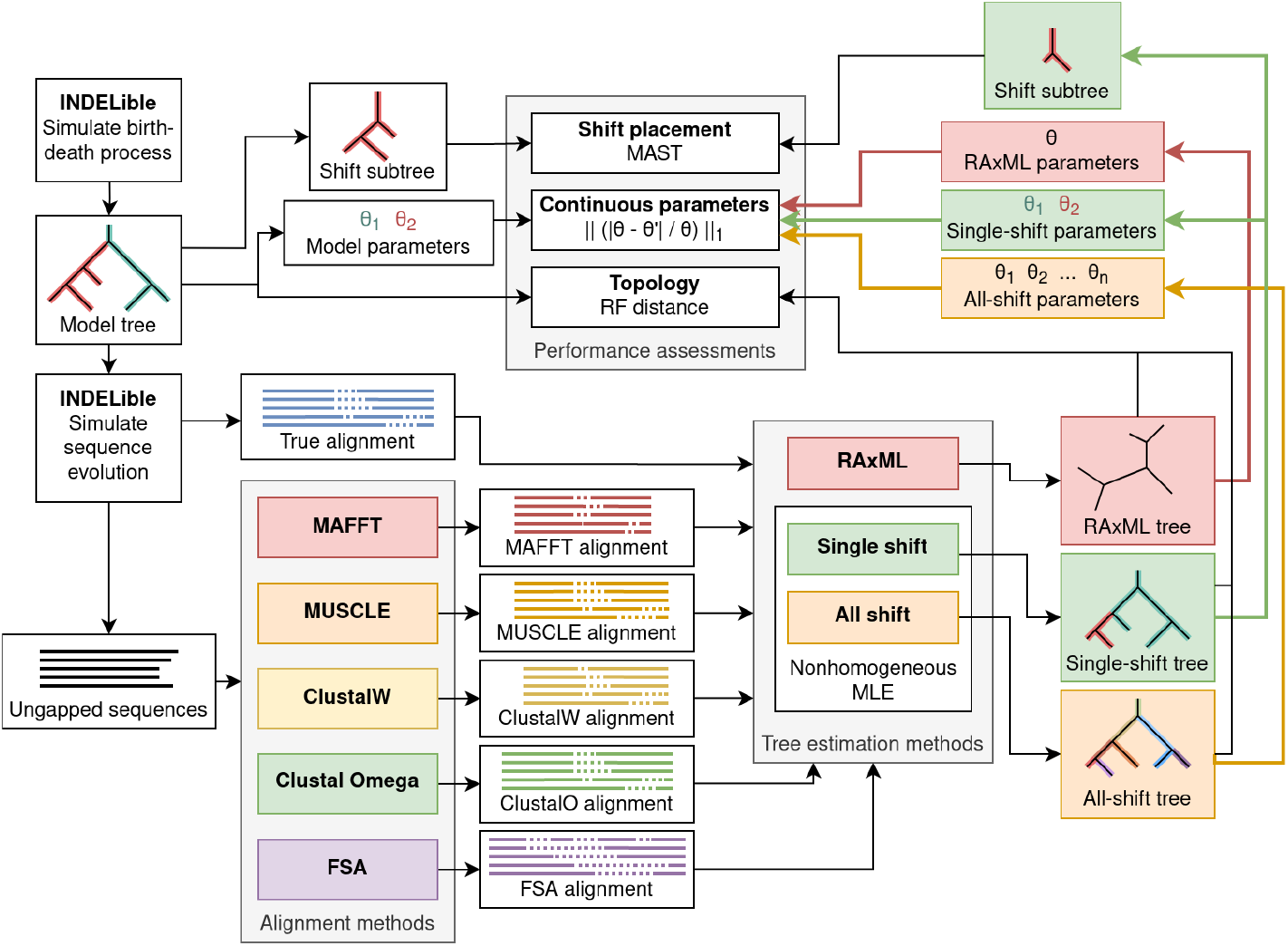
Flowchart of simulation study steps. A model tree is first generated under a birth-death process using INDELible, and then sequences are generated under that model tree with a nonhomogeneous model. The ungapped sequences are aligned using MAFFT, MUSCLE, ClustalW, Clustal Omega, and FSA. Every estimated alignment, as well as the true alignment, is then used to perform tree estimation. Tree estimation is done using three different classes of model: 0-shift, or homogeneous, is done using RAxML. Single-shift and all-shift, both nonhomogeneous across lineages, are performed using PAML to calculate likelihood scores and continuous parameter optimization. Finally, the resulting trees from each alignment and tree estimation method pair is compared against the model tree. For the single shift model, shift placement is evaluated by computing the size of the maximum agreement subtree (MAST) for the true and estimated shift subtrees.

#### Simulating sequence evolution

Model conditions were the same as those used in **?** to include a range of sequence divergence in the simulation study. INDELible was used to generate sequences under a GTR-based branch model using the phylogenies generated as described earlier. **?** found GC content variation in the avian phylogeny. The GTR model parameters were empirically estimated using single-copy orthologs from **?** for the subset of species (*Calypte anna, Alligator mississippiensis, Melopsittacus undulatus, Corvus brachyrhynchos*, and *Manacus vitellinus*) included in **?**’s study. To estimate these parameters, we aligned the single-copy orthologs using MAFFT with the default settings. Then, we used a single-shift model to estimate parameters on each individual aligned sequence. Then, we looked at the two sets of substitution rates estimated, and observed that the ratio between them was bimodal. The first peak ranged from a 1- to 10-fold difference, and the second ranged from a 10000- to 100000-fold difference. We chose GTR model parameters based on the estimated parameters in the first mode.

### 2.3 Empirical study

#### Grass dataset

The distribution of GC content in monocots is bimodal [**?**], which is not the case for other plants. This pattern is notably strong in rice. We applied nonhomogeneous substitution model based phylogenetic tree estimation to a set of 8 taxa: Oryza sativa japonica [**?**], Sorghum bicolor [**?**], Carex cristatella, Carex scoparia, Juncus effusus, Juncus inflexus [**?**], Ananas comosus [**?**], and Musa balbisiana [**?**]. We identified 1900 single-copy orthologs using orthofinder [**?**] with the default settings. We aligned the sequences individually using MAFFT, MUSCLE, Clustal Omega, Clustal W, and FSA with the default settings. We performed phylogenetic estimation under a single-shift model for every individual gene. We also concatenated the aligned gene sequences and ran the same analysis.

## 3 Results

### 3.1 Simulation study

#### Impact of alignment accuracy

In both topology estimation and substitution model parameter estimation, the true alignments perform the best, as would be expected. Across all methods and levels of sequence divergence, topological inference using estimated alignments yields significantly more error. In figure **??**, we see a correlation between topological error and alignment accuracy in more divergent model conditions. We also see this trend is maintained in the 20-taxa model conditions.

#### Impact of model misspecification

Nonstationary nucleotide composition and nonhomogeneous substitution rates can reflect an evolutionary adaptation. Estimates for base frequencies and substitution rates can be used to characterize such adaptations. For substitution model parameter estimation, using the single-shift model yields the closest parameter estimates across all alignment types. In figure **??** we can see that for topological estimation, depending on the alignment and level of divergence, the single-shift model performs as well as, but usually better than the underspecified model used with RAxML. The overspecified model performs about as well as well as the homogeneous model until the 20E model condition.

**Table 2:** Model conditions and summary statistics for ground truth and estimated alignments.

| Model condition | # Taxa | Tree height | Indel probability | Length | ANHD | Gappiness | Alignment | SP-FP | SP-FN | Estimated length | Estimated NHD | Estimated gappiness |
| --- | --- | --- | --- | --- | --- | --- | --- | --- | --- | --- | --- | --- |
| 10.A | 10 | 0.47 | 0.13 | 2123.8 | 0.306 | 0.528 | MAFFT | 0.572 | 0.512 | 1478.6 | 0.389 | 0.326 |
|  |  |  |  |  |  |  | MUSCLE | 0.579 | 0.508 | 1518.5 | 0.413 | 0.342 |
|  |  |  |  |  |  |  | CLUSTALW | 0.746 | 0.683 | 1191.4 | 0.466 | 0.165 |
|  |  |  |  |  |  |  | CLUSTALO | 0.734 | 0.682 | 1247.1 | 0.484 | 0.202 |
|  |  |  |  |  |  |  | FSA | 0.217 | 0.645 | 3609.3 | 0.280 | 0.715 |
| 10.B | 10 | 0.7 | 0.1 | 2315.8 | 0.364 | 0.564 | MAFFT | 0.683 | 0.629 | 1477.1 | 0.435 | 0.321 |
|  |  |  |  |  |  |  | MUSCLE | 0.667 | 0.602 | 1570.1 | 0.449 | 0.361 |
|  |  |  |  |  |  |  | CLUSTALW | 0.786 | 0.724 | 1186.0 | 0.496 | 0.155 |
|  |  |  |  |  |  |  | CLUSTALO | 0.781 | 0.732 | 1248.7 | 0.504 | 0.198 |
|  |  |  |  |  |  |  | FSA | 0.236 | 0.603 | 4471.2 | 0.319 | 0.770 |
| 10.C | 10 | 1.2 | 0.06 | 2313.2 | 0.465 | 0.566 | MAFFT | 0.752 | 0.711 | 1484.5 | 0.484 | 0.328 |
|  |  |  |  |  |  |  | MUSCLE | 0.729 | 0.679 | 1573.1 | 0.499 | 0.364 |
|  |  |  |  |  |  |  | CLUSTALW | 0.822 | 0.768 | 1170.6 | 0.537 | 0.148 |
|  |  |  |  |  |  |  | CLUSTALO | 0.823 | 0.780 | 1237.6 | 0.541 | 0.194 |
|  |  |  |  |  |  |  | FSA | 0.272 | 0.783 | 4992.4 | 0.377 | 0.795 |
| 10.D | 10 | 2 | 0.031 | 2202.8 | 0.553 | 0.538 | MAFFT | 0.828 | 0.807 | 1461.8 | 0.528 | 0.310 |
|  |  |  |  |  |  |  | MUSCLE | 0.794 | 0.766 | 1561.9 | 0.542 | 0.353 |
|  |  |  |  |  |  |  | CLUSTALW | 0.865 | 0.830 | 1143.0 | 0.573 | 0.119 |
|  |  |  |  |  |  |  | CLUSTALO | 0.858 | 0.831 | 1222.9 | 0.565 | 0.177 |
|  |  |  |  |  |  |  | FSA | 0.384 | 0.864 | 5729.6 | 0.420 | 0.820 |
| 10.E | 10 | 4.4 | 0.013 | 2063.0 | 0.649 | 0.510 | MAFFT | 0.879 | 0.871 | 1529.2 | 0.569 | 0.342 |
|  |  |  |  |  |  |  | MUSCLE | 0.846 | 0.831 | 1590.5 | 0.589 | 0.366 |
|  |  |  |  |  |  |  | CLUSTALW | 0.897 | 0.872 | 1146.5 | 0.614 | 0.124 |
|  |  |  |  |  |  |  | CLUSTALO | 0.884 | 0.865 | 1234.0 | 0.602 | 0.187 |
|  |  |  |  |  |  |  | FSA | 0.497 | 0.912 | 6196.2 | 0.477 | 0.836 |
| 20.A | 20 | 0.47 | 0.13 | 2410.3 | 0.301 | 0.581 | MAFFT | 0.420 | 0.388 | 1643.7 | 0.368 | 0.390 |
|  |  |  |  |  |  |  | MUSCLE | 0.404 | 0.351 | 1790.2 | 0.380 | 0.439 |
|  |  |  |  |  |  |  | CLUSTALW | 0.654 | 0.612 | 1278.6 | 0.449 | 0.216 |
|  |  |  |  |  |  |  | CLUSTALO | 0.654 | 0.625 | 1306.3 | 0.471 | 0.233 |
|  |  |  |  |  |  |  | FSA | 0.129 | 0.536 | 4394.5 | 0.289 | 0.763 |
| 20.B | 20 | 0.7 | 0.1 | 2585.1 | 0.374 | 0.606 | MAFFT | 0.554 | 0.526 | 1670.7 | 0.433 | 0.394 |
|  |  |  |  |  |  |  | MUSCLE | 0.521 | 0.470 | 1863.0 | 0.445 | 0.456 |
|  |  |  |  |  |  |  | CLUSTALW | 0.753 | 0.713 | 1261.2 | 0.509 | 0.198 |
|  |  |  |  |  |  |  | CLUSTALO | 0.738 | 0.708 | 1318.2 | 0.518 | 0.232 |
|  |  |  |  |  |  |  | FSA | 0.144 | 0.681 | 5959.0 | 0.349 | 0.819 |
| 20.C | 20 | 1.2 | 0.06 | 2895.8 | 0.484 | 0.649 | MAFFT | 0.771 | 0.752 | 1683.0 | 0.516 | 0.400 |
|  |  |  |  |  |  |  | MUSCLE | 0.720 | 0.684 | 1907.7 | 0.530 | 0.470 |
|  |  |  |  |  |  |  | CLUSTALW | 0.856 | 0.822 | 1227.8 | 0.573 | 0.178 |
|  |  |  |  |  |  |  | CLUSTALO | 0.848 | 0.821 | 1303.2 | 0.570 | 0.226 |
|  |  |  |  |  |  |  | FSA | 0.215 | 0.838 | 8231.8 | 0.429 | 0.874 |
| 20.D | 20 | 2 | 0.031 | 2696.2 | 0.581 | 0.622 | MAFFT | 0.863 | 0.860 | 1691.4 | 0.569 | 0.403 |
|  |  |  |  |  |  |  | MUSCLE | 0.815 | 0.800 | 1890.0 | 0.585 | 0.466 |
|  |  |  |  |  |  |  | CLUSTALW | 0.897 | 0.876 | 1181.6 | 0.615 | 0.148 |
|  |  |  |  |  |  |  | CLUSTALO | 0.890 | 0.875 | 1287.4 | 0.602 | 0.218 |
|  |  |  |  |  |  |  | FSA | 0.328 | 0.908 | 10177.9 | 0.486 | 0.899 |
| 20.E | 20 | 4.4 | 0.013 | 2723.6 | 0.667 | 0.629 | MAFFT | 0.939 | 0.940 | 1804.3 | 0.608 | 0.444 |
|  |  |  |  |  |  |  | MUSCLE | 0.904 | 0.898 | 1979.5 | 0.630 | 0.493 |
|  |  |  |  |  |  |  | CLUSTALW | 0.946 | 0.933 | 1162.8 | 0.652 | 0.140 |
|  |  |  |  |  |  |  | CLUSTALO | 0.938 | 0.928 | 1283.8 | 0.637 | 0.221 |
|  |  |  |  |  |  |  | FSA | 0.432 | 0.942 | 11566.5 | 0.525 | 0.913 |

### 3.2 Empirical study

#### Grass dataset

For the concatenated analysis, the estimated topology was identical for all alignment types except ClustalW. Furthermore, the placement of the shift was identical for all alignment types except ClustalW. The estimated tree using a no homogeneous model with the concatenated MAFFT alignment identifies Ananas comosus as more closely related to Oryza sativa than Sorghum bicolor, which would be a very unconvential result. The tree estimated using a homogeneous model on the MAFFT alignment does not make this placement, and is in consensus with Clustal Omega, ClustalW, and Muscle estimated tree topologies.

**Figure 2:**
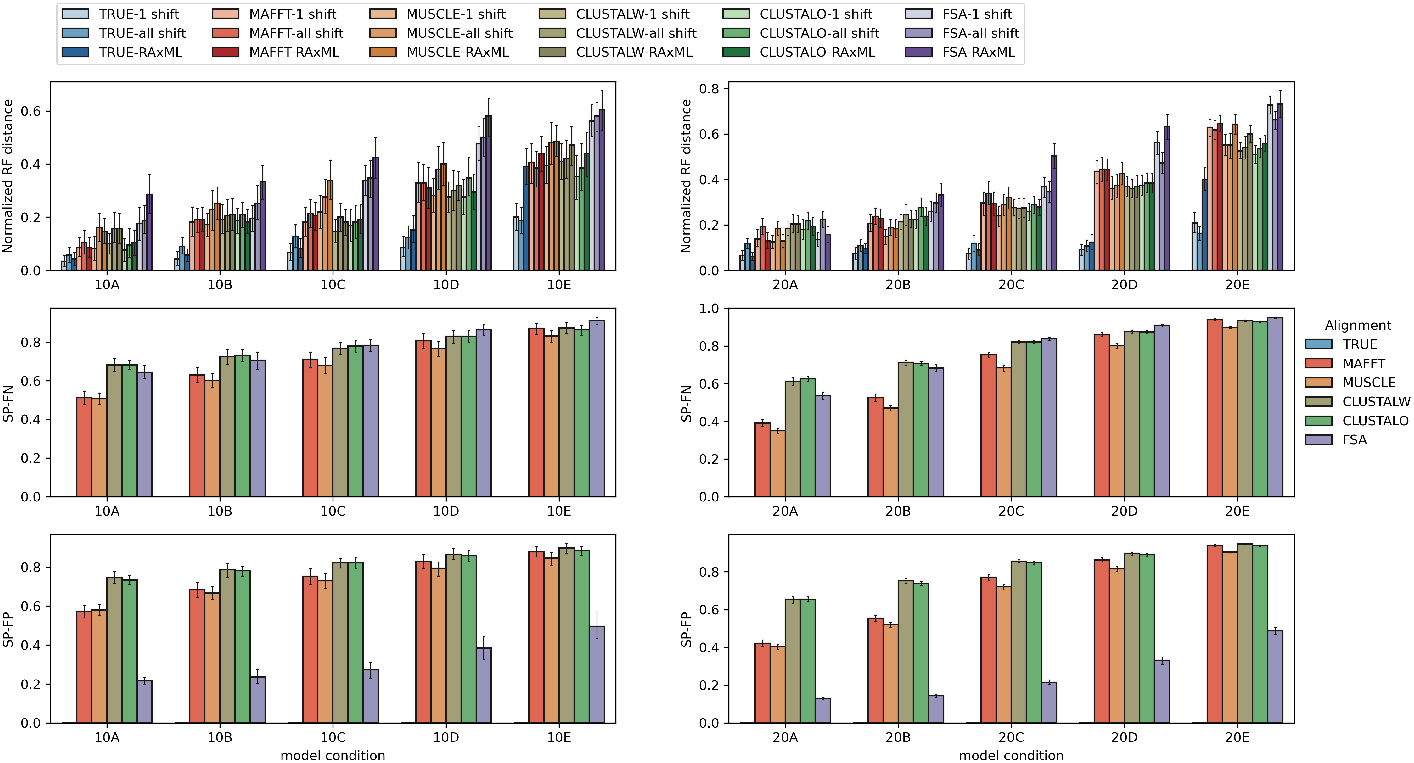
Topological error for every combination of alignment method and maximum likelihood tree estimation method. Topological error is measured with the normalized Robinson-Foulds distance of the model tree and the tree estimated using a single-shift nonhomogeneous substitution model. Alignment SP-FN and SP-FP are also reported.

**Figure 3:**
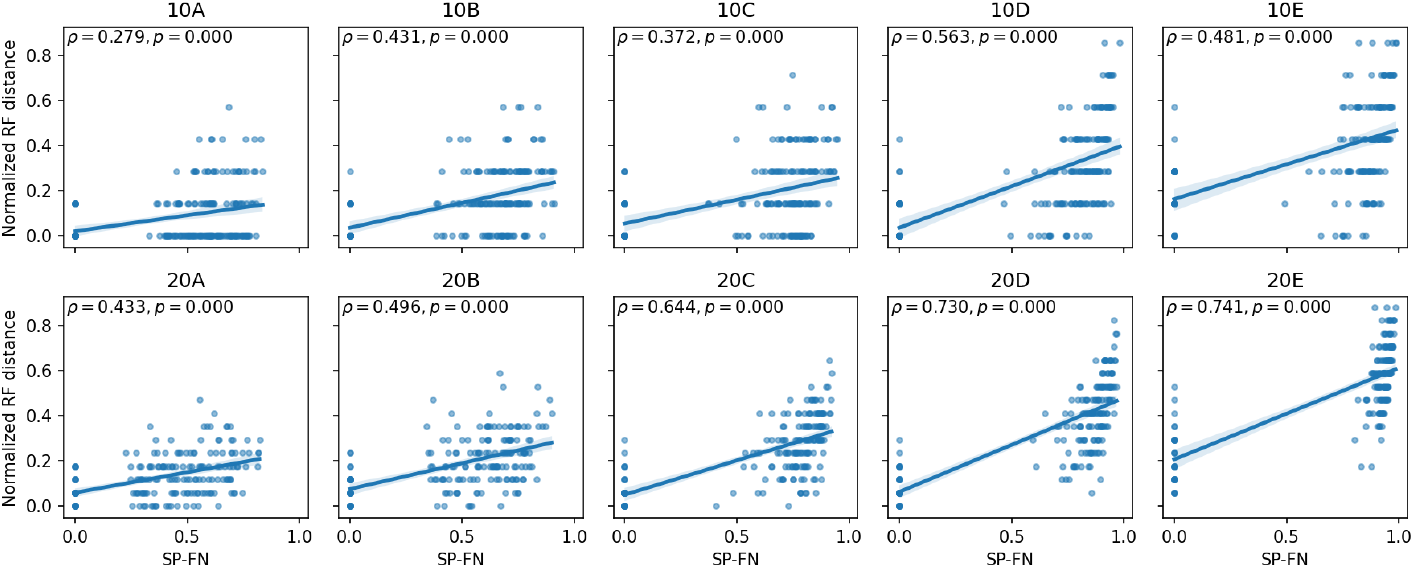
Alignment error (sum-of-pairs false negative rate) vs topological error (normalized Robinson-Foulds distance). Results are aggregated for all alignment types.

**Figure 4:**
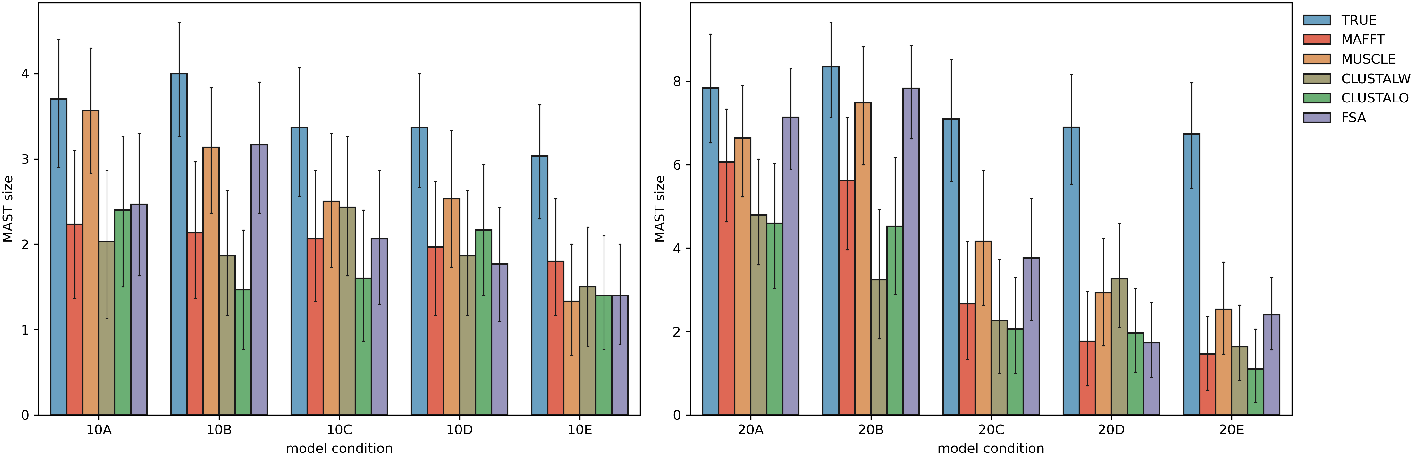
Size of the MAST of the predicted shift subtree and the true subtree.

**Table 3:**
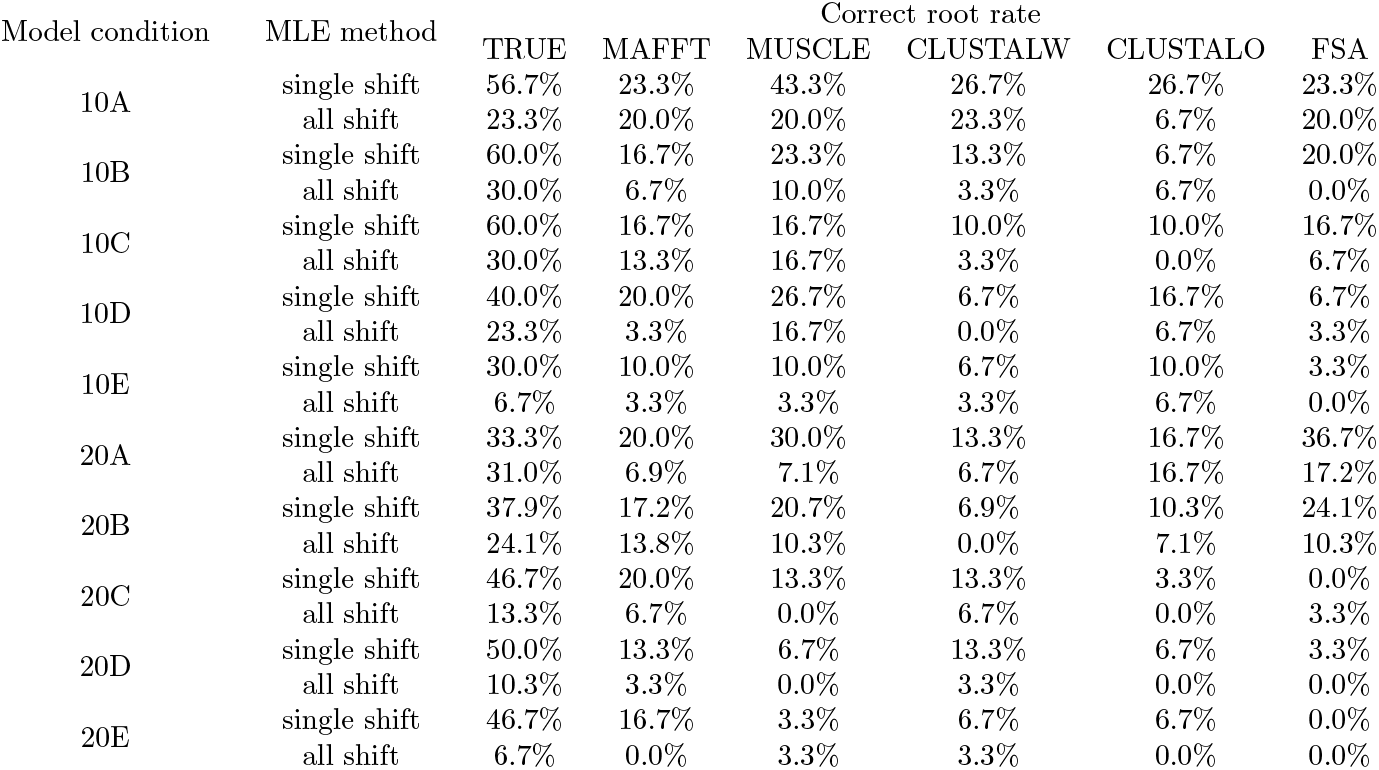
Proportion of correct root placements by alignment and model condition.

| Model condition | MLE method | Correct root rate |  |  |  |  |  |
| --- | --- | --- | --- | --- | --- | --- | --- |
|  |  | TRUE | MAFFT | MUSCLE | CLUSTALW | CLUSTALO | FSA |
| 10A | single shift | 56.7% | 23.3% | 43.3% | 26.7% | 26.7% | 23.3% |
|  | all shift | 23.3% | 20.0% | 20.0% | 23.3% | 6.7% | 20.0% |
| 10B | single shift | 60.0% | 16.7% | 23.3% | 13.3% | 6.7% | 20.0% |
|  | all shift | 30.0% | 6.7% | 10.0% | 3.3% | 6.7% | 0.0% |
| 10C | single shift | 60.0% | 16.7% | 16.7% | 10.0% | 10.0% | 16.7% |
|  | all shift | 30.0% | 13.3% | 16.7% | 3.3% | 0.0% | 6.7% |
| 10D | single shift | 40.0% | 20.0% | 26.7% | 6.7% | 16.7% | 6.7% |
|  | all shift | 23.3% | 3.3% | 16.7% | 0.0% | 6.7% | 3.3% |
| 10E | single shift | 30.0% | 10.0% | 10.0% | 6.7% | 10.0% | 3.3% |
|  | all shift | 6.7% | 3.3% | 3.3% | 3.3% | 6.7% | 0.0% |
| 20A | single shift | 33.3% | 20.0% | 30.0% | 13.3% | 16.7% | 36.7% |
|  | all shift | 31.0% | 6.9% | 7.1% | 6.7% | 16.7% | 17.2% |
| 20B | single shift | 37.9% | 17.2% | 20.7% | 6.9% | 10.3% | 24.1% |
|  | all shift | 24.1% | 13.8% | 10.3% | 0.0% | 7.1% | 10.3% |
| 20C | single shift | 46.7% | 20.0% | 13.3% | 13.3% | 3.3% | 0.0% |
|  | all shift | 13.3% | 6.7% | 0.0% | 6.7% | 0.0% | 3.3% |
| 20D | single shift | 50.0% | 13.3% | 6.7% | 13.3% | 6.7% | 3.3% |
|  | all shift | 10.3% | 3.3% | 0.0% | 3.3% | 0.0% | 0.0% |
| 20E | single shift | 46.7% | 16.7% | 3.3% | 6.7% | 6.7% | 0.0% |
|  | all shift | 6.7% | 0.0% | 3.3% | 3.3% | 0.0% | 0.0% |

For the per-gene analyses, figure **??** shows that on average, the topologies estimated from different alignment methods have some disagreement. For continuous parameter estimates table **??** shows that for all alignments, estimated shift and background models usually showed some difference in base frequencies and substitution rate estimates.

Figure **??** shows that the topology estimated was identical across alignments. The rooting for ClustalW was different, as was the shift placement.

## 4 Discussion

### Alignment quality

Our results from the simulation study supports that alignment accuracy can have an impact on tree topology estimation. Substitution models accounting for nonhomogeneous and nonstationary sequence evolution can be used to study the origins and consequences of substitution rate variation across lineages. From both the simulation and empirical study, downstream estimates of substitution rates can vary between different alignment methods.

**Figure 5:**
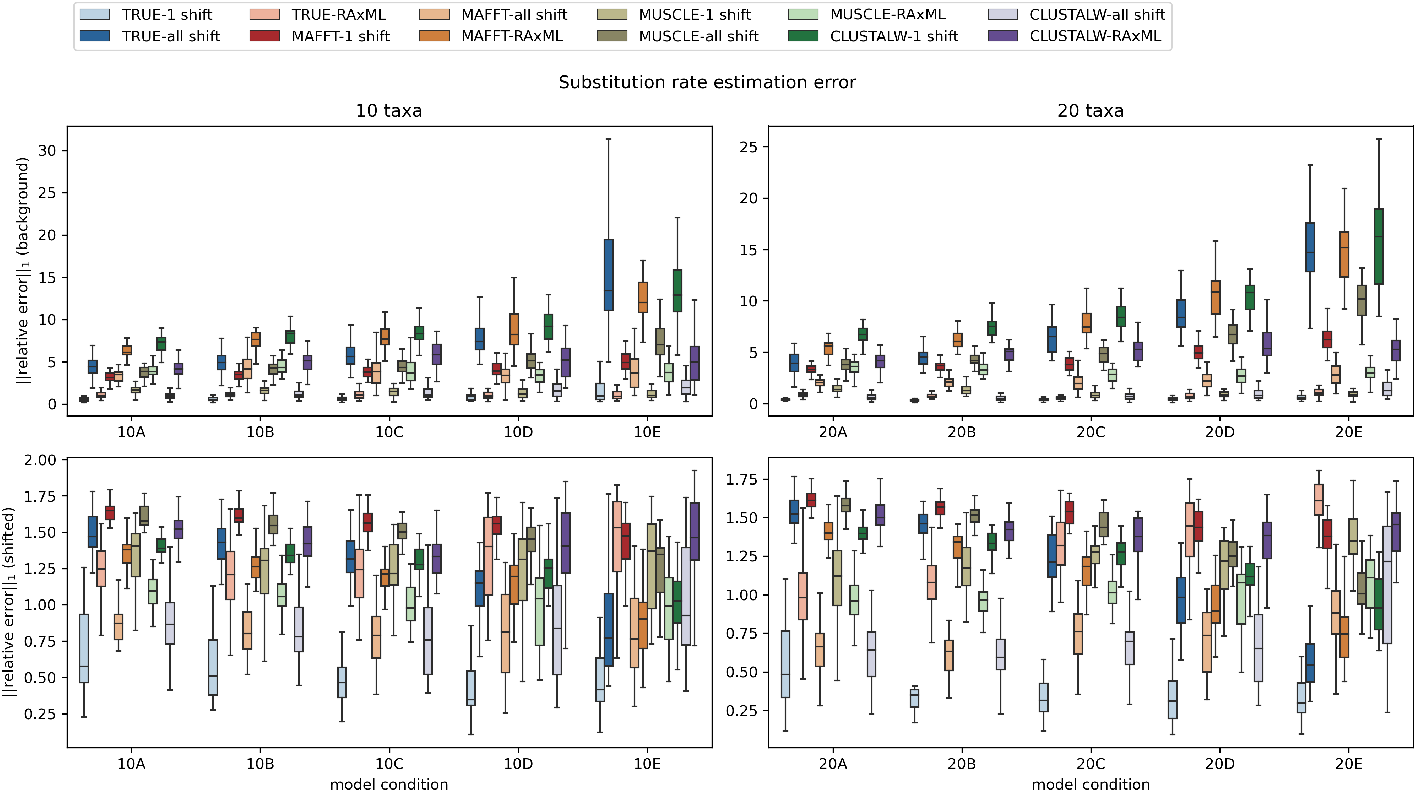
Substitution rate estimation error as measured by L1-norm of the relative errors for each exchangeability parameter.

**Figure 6:**
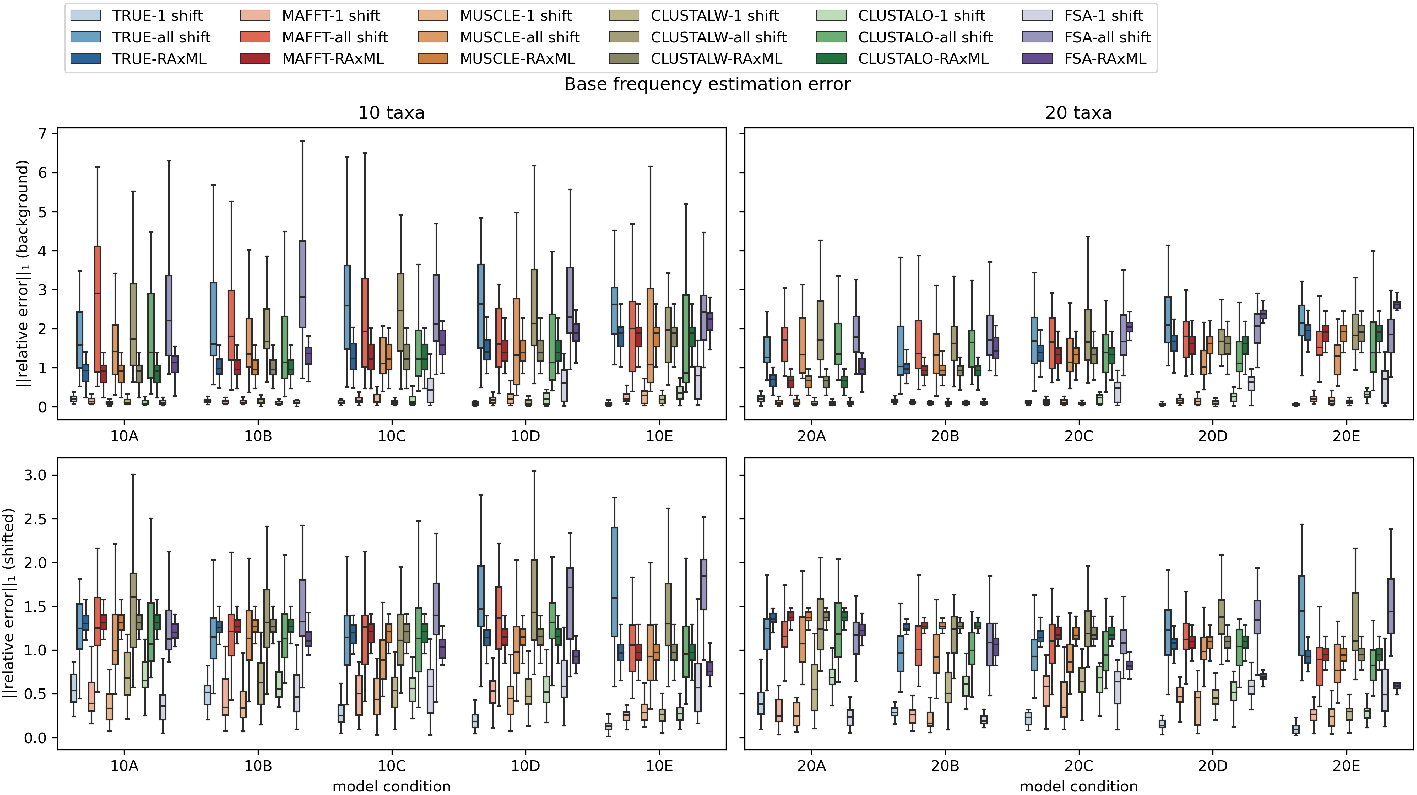
Base frequency estimation error as measured by the L1 norm of the relative errors for each base frequency.

FSA exhibits the most underalignment, as shown in table **??**. This seems to negatively impact tree topology estimation far more than substitution model parameter estimation. Conversely, MUSCLE and Clustal Omega perform relatively well in the task of topology estimation among the selected alignment methods, but poorly estimate background model parameters. However, there isn’t an obvious pattern from the alignment summary statistics that would point to some quality of the alignments resulting from these methods that leads to this difference.

**Table 4:** Aggregated statistics of Robinson-Foulds distance of gene trees estimated using a single-shift model.

|  | muscle |  | clustalo |  | clustalw |  | fsa |  |
| --- | --- | --- | --- | --- | --- | --- | --- | --- |
| nRF | Mean | Std | Mean | Std | Mean | Std | Mean | Std |
| mafft | 0.044 | 0.088 | 0.052 | 0.097 | 0.059 | 0.106 | 0.041 | 0.087 |
| muscle |  |  | 0.053 | 0.100 | 0.058 | 0.108 | 0.044 | 0.090 |
| clustalo |  |  |  |  | 0.059 | 0.105 | 0.052 | 0.095 |
| clustalw |  |  |  |  |  |  | 0.061 | 0.103 |
| nRF (rooted) | Mean | Std | Mean | Std | Mean | Std | Mean | Std |
| mafft | 0.171 | 0.191 | 0.188 | 0.200 | 0.191 | 0.198 | 0.191 | 0.205 |
| muscle |  |  | 0.192 | 0.204 | 0.191 | 0.203 | 0.198 | 0.206 |
| clustalo |  |  |  |  | 0.192 | 0.204 | 0.213 | 0.205 |
| clustalw |  |  |  |  |  |  | 0.217 | 0.207 |

**Figure 7:**
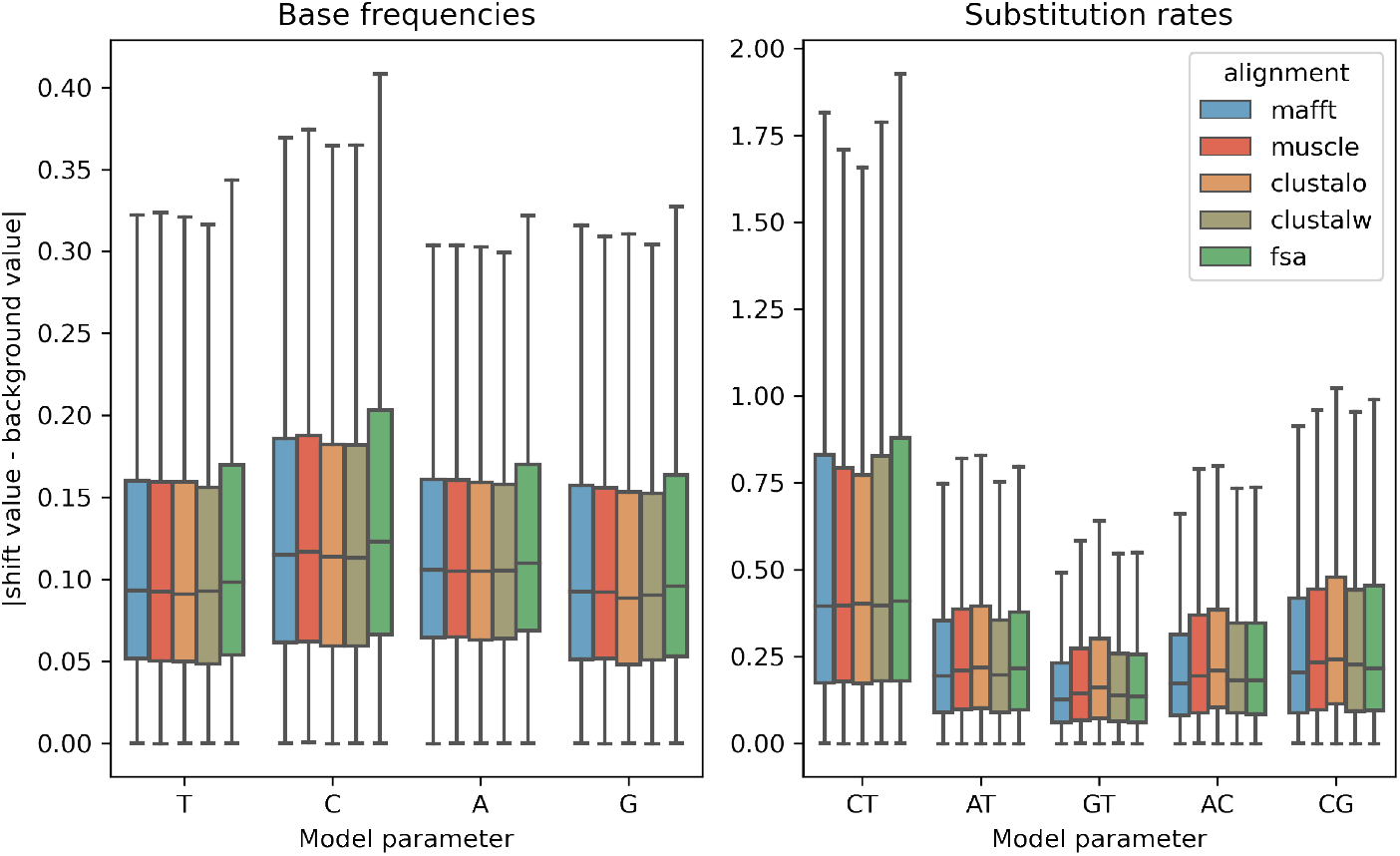
Box plot of the difference between background and shift model parameters estimated for each of the single copy orthologs. Median and interquartile range is represented, whiskers are 1.5IQR and values outside of the whiskers are not shown.

**Figure 8:**
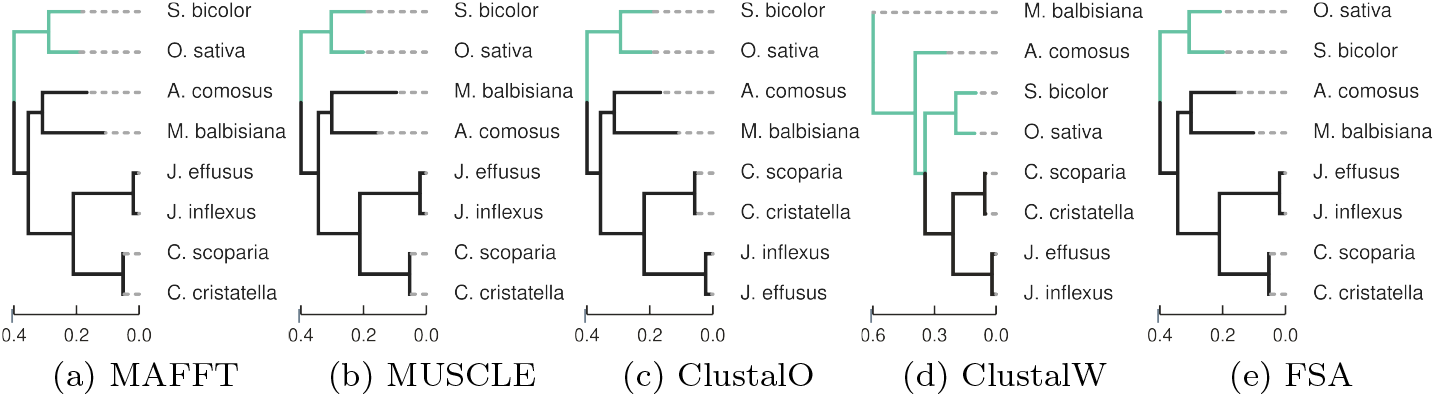
Tree topologies for the grass dataset estimated using concatenated MAFFT, MUSCLE, Clustal Omega, ClustalW, and FSA alignments respectively. Branches that are predicted to have evolved with elevated G+C base frequencies relative to the rest of the tree are highlighted in green.

**Table 5:** Model parameter estimates from the concatenated analysis

| Alignment | Which model | T | C | A | G | C↔ T | A↔ T | G↔ T | A↔ C | C↔ G |
| --- | --- | --- | --- | --- | --- | --- | --- | --- | --- | --- |
| mafft | shift | 0.294 | 0.176 | 0.313 | 0.217 | 1.245 | 0.344 | 0.336 | 0.511 | 0.600 |
| mafft | background | 0.234 | 0.254 | 0.243 | 0.269 | 1.087 | 0.420 | 0.313 | 0.432 | 0.585 |
| muscle | shift | 0.294 | 0.178 | 0.311 | 0.217 | 1.233 | 0.389 | 0.380 | 0.580 | 0.676 |
| muscle | background | 0.234 | 0.254 | 0.243 | 0.269 | 1.082 | 0.450 | 0.349 | 0.482 | 0.675 |
| clustalo | shift | 0.295 | 0.179 | 0.309 | 0.218 | 1.194 | 0.416 | 0.406 | 0.627 | 0.722 |
| clustalo | background | 0.234 | 0.253 | 0.243 | 0.269 | 1.071 | 0.460 | 0.359 | 0.503 | 0.705 |
| clustalw | shift | 0.287 | 0.169 | 0.331 | 0.213 | 1.343 | 0.389 | 0.410 | 0.585 | 0.615 |
| clustalw | background | 0.246 | 0.241 | 0.251 | 0.262 | 1.111 | 0.450 | 0.341 | 0.479 | 0.575 |
| fsa | shift | 0.291 | 0.178 | 0.316 | 0.215 | 1.251 | 0.350 | 0.341 | 0.513 | 0.605 |
| fsa | background | 0.231 | 0.259 | 0.242 | 0.268 | 1.075 | 0.427 | 0.314 | 0.427 | 0.579 |

In the concatenated empirical study, there was consensus between all MSA methods except ClustalW topologically as well as for shift placement. Even when averaged across all loci, substitution rate estimates were highly variable between alignment methods. The simulation study results suggest that it is more difficult to estimate model parameters for the more basal substitution model. From the standard deviation columns in **??**, we see there’s less variance across model parameter estimates for the model with a higher estimated G+C base frequencies across loci. The model with elevated G+C typically corresponds to the subtree containing Oryza sativa and Sorghum bicolor.

### Branch model of substitution

In the simulation study, phylogenetic estimation using a branch model matching the number of shifts that the sequence data was evolved under gives the best performance, as expected. Interestingly, the all-shift model performs very closely to the no-shift model for topology estimation, except in the 20E model condition, where it performs slightly better than the no-shift model. This difference in performance may indicate that the improvement of these models is greater when there are more sequences being studied and they are more divergent. Though in most cases this seems to suggest that the most general overparameterization doesn’t provide a significant improvement in topological estimation over model misspecification.

A generalization of single-shift branch model, which we’ll call k-shift, can be described with the aim to achieve the simplest explanation. This model would make the tradeoff of having less continuous parameter estimation in exchange for having a larger search space to explore, but signals of strong shifts might be useful for narrowing this search space.

### Limitations

This study looks at a very specific case where exactly one change has occurred in the phylogeny. We demonstrate that even in this simplified case, all aspects of phylogenetic estimation are impacted by both alignment quality and model misspecification. In this scenario, we assumed that there were exactly two sets of substitution model parameters, and that one of those sets applied to all the descending edges from a starting edge. As it is, the search space of branch model assignments for each tree topology is *O*(*n*) where *n* is the number of taxa. Natural extensions would be to allow for more sets of substitution model parameters as well as less restrictions on what assignment of these models to the branches are considered. These relaxations drastically increase the size of the search space for model assignments to the tree topology, as well as the number of continuous parameters to optimize for with the former. Because of this, this study did not look at how a *k*-shift model would perform in the case that there were potentially more sets of substitution model parameters.

A *k*-shift model is a natural extension, but would pose several challenges for estimation on empirical data as well. Furthermore, these would also limit our ability to look at realistic model conditions. One issue is in the availability of empirical data with novel observations of multiple compositional shifts to gather estimates from. Most studies only make note of two categories, usually high GC and low GC. Another arises from how quickly the search space increases, as finding multiple significant changes in internal branches would necessitate more taxa to study.

## 5 Conclusions

In both the simulation and empirical studies, we looked at how both alignment accuracy and model misspecification had an impact on downstream phylogenetic inference and estimation. In our simulation study, we looked at a scenario where there’s exactly one change in the substitution model that occurs in the simulated tree, and that it occurs such that all descending edges also evolve with that model. We showed that even in this simple case, MSA quality affected all aspects of downstream phylogenetic estimation using a nonhomogeneous model, from tree topology to continuous parameter estimation. Furthermore, we found using a nonhomogeneous substitution model for maximum likelihood estimation yielded closer to ground truth results than using a homogeneous substitution model.

In our empirical study, we observed an impact in tree topology estimation when using a nonhomogeneous model versus a homogeneous substitution model, supporting that the homogeneity-across-lineages assumption can affect estimation even when dealing with large concatenated alignments. Furthermore, we found that estimates using different alignments had a fair amount of disagreement between their estimated gene tree topologies, and estimated continuous parameters were even more sensitive to the alignment method used.

## Acknowledgements

We thank Kevin Childs and Yu-ya Liang for help with the grass dataset. This work is supported in part by the National Science Foundation Research Traineeship Program (DGE-1828149) to Rei Doko and through computational resources and services provided by the Institute for Cyber-Enabled Research at Michigan State University.

