## Supplementary Online Materials for "The Impact of Multiple Sequence Alignment Error on Phylogenetic Estimation under Variable-Across-Phylogeny Substitution Models"

#### 1 Supplementary Methods

**Software commands used.** INDELible [Fletcher and Yang, 2009] version 1.03 was run using the following settings to simulate model tree evolution

```
[TYPE] NUCLEOTIDE 1
[MODEL] mymodel
    [submodel] JC
[TREE] mytree
    [rooted] <# taxa>
[PARTITIONS] mypartition [mytree mymodel 1]
[EVOLVE] mypartition <# replicates> output
```

The trees are rescaled to have nonultrametric branch lengths and the following control settings to simulate sequence evolution:

```
[TYPE] NUCLEOTIDE 1
[SETTINGS]
  [output] PHYLIP
[MODEL] background
  [submodel] GTR <CT> <AT> <GT> <AC> <CG>
  [statefreq] <T> <C> <A> <G>
  [indelmodel] USER <path to indel distribution>
  [indelrate] <indel rate>
[MODEL] shift
  [submodel] GTR <CT> <AT> <GT> <AC> <CG>
  [statefreq] <T> <C> <A> <G>
  [indelmodel] USER <path to indel distribution>
  [indelrate] <indel rate>
[TREE] mytree <tree>
  [treedepth] <specified tree height>
[BRANCHES] mymodel <tree with model placements defined>
[PARTITIONS] mypartition [mytree mymodel <sequence length>]
[EVOLVE] mypartition 1 sequence
```

The following command was used to perform MSA estimation with MAFFT [Katoh and Standley, 2013] version 7.475

```
mafft <input sequence> <output alignment>
```

MUSCLE [Edgar, 2004] version 5.0.1428 was run with the following:

```
muscle -align <input sequence> -output <output alignment>
```

Clustal Omega [Sievers et al., 2011] version 1.2.4 was run with the following

```
clustalo -i <input sequence> -t DNA --threads 1 ><output alignment>
```

ClustalW [Larkin et al., 2007] version 2.1 was run with the following

```
clustalw2 <input sequence> -type=DNA -outfile=<output alignment>
```

FSA [Sievers et al., 2011] version 1.15.9 was run with the following

```
fsa <input sequence --maxram 8192 ><output alignment>
```

For single-shift search, PAML [Yang, 2007] was run with the following control file:

```
     seqfile = <sequence_path>
    treefile = <tree path>
     outfile = <result output path>
       noisy = 3
     verbose = 3
     runmode = 0
       model = 7   * GTR model
       Mgene = 0
       ndata = 1
       nhomo = 5
   fix_kappa = 2
       clock = 0
   fix_alpha = 1
       alpha = 0.
       getSE = 0
RateAncestor = 0
   cleandata = 0
      method = 0
 fix_blength = 0
```

The control file for the all-shift model is identical, except:

```
       nhomo = 3
   fix_kappa = 0
```

RAxML [Stamatakis, 2014] was run with the following command:

```
raxml -s <msa path> -n <name> -m GTRCAT -V -p <random number>
```

**Grass dataset processing.** To obtain single copy orthologs for the grass dataset, we ran orthofinder using the following command:

```
orthofinder -f <path containing sequence files>
```

### 2 Supplementary Results and Discussion.

**Simulation study runtime and memory usage.** Memory usage for the nonhomogeneous substitution model did not exceed 1 GB.

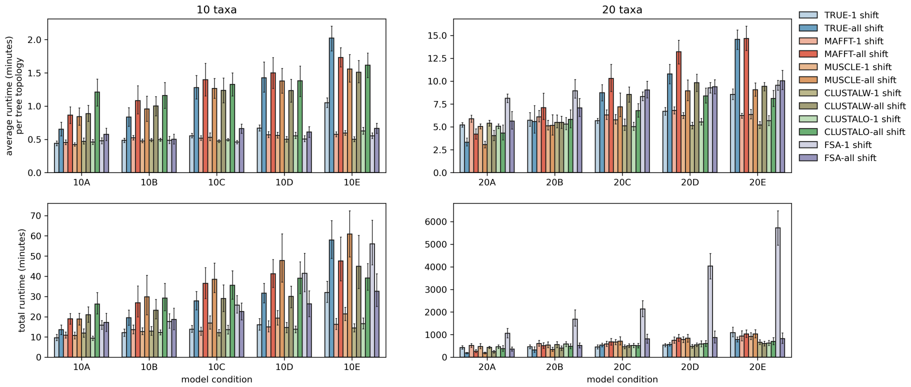


Figure 1: Runtime for nonhomogeneous tree search. The first row represents how long it takes for PAML to do continuous parameter optimization and likelihood calculation, and the second row represents the total time the wrapper script takes.

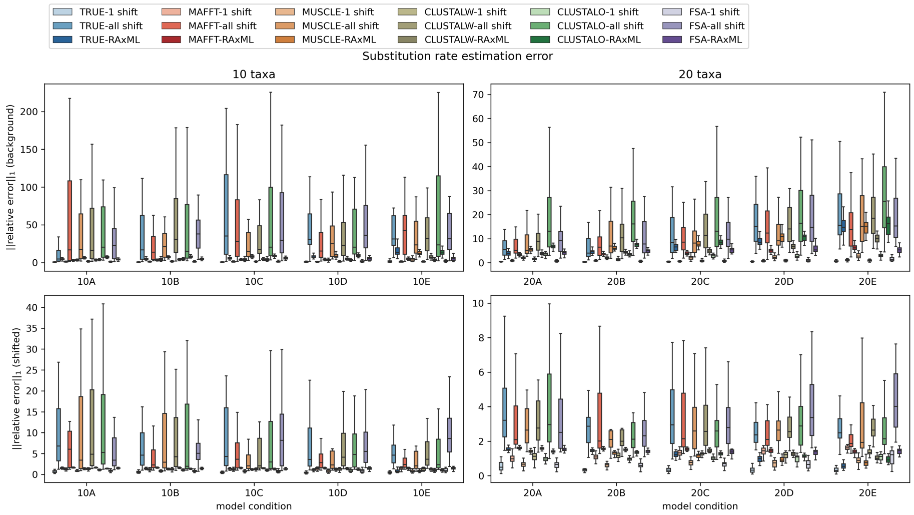


Figure 2: Substitution rate error.

**Estimation error** For branch lengths, there wasn't a clean way to quantify error, in part because tree estimation and branch length estimation are heavily intertwined. Comparing leaf-edge distances only captures roughly half of the estimated branch lengths, but comparing pairwise distances does not consider the topology in any way. Kuhner-Felsenstein distance [Kuhner and Felsenstein, 1994], shown in the middle panels in 3, takes topology into consideration, but is difficult to interpret. For this reason we cannot draw any meaningful conclusions about branch length estimation.

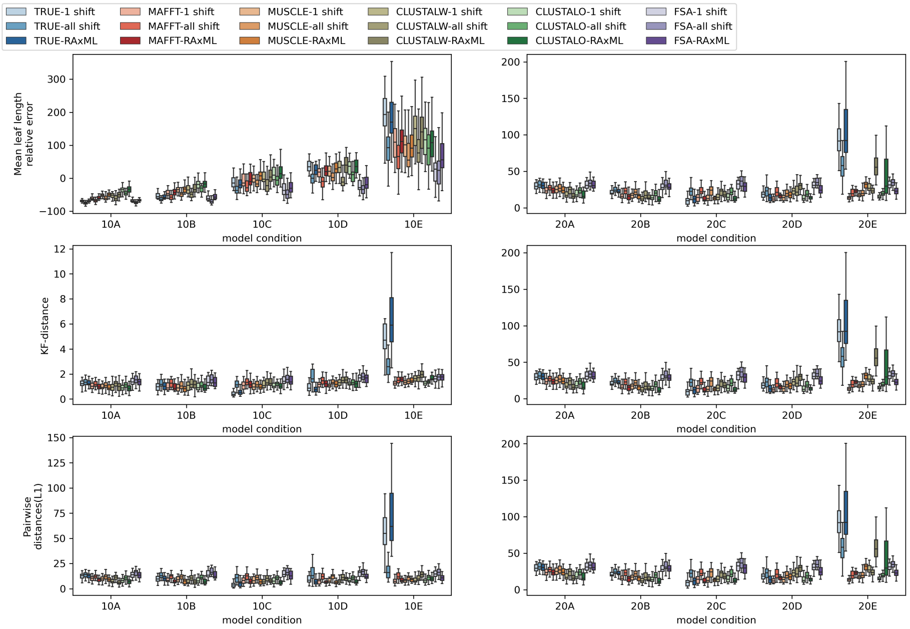


Figure 3: Various measures for branch length estimation error. KF stands for Kuhner and Felsenstein [1994].
